# Mathematical Modelling of Microtube-Driven Regrowth of Glioma After Local Resection

**DOI:** 10.1101/2024.08.20.608880

**Authors:** Alexandra Shyntar, Thomas Hillen

**Affiliations:** University of Alberta

**Keywords:** tumor microtubes, glioblastoma modelling, anisotropic diffusion

## Abstract

Recently, glioblastoma tumors were shown to form tumor microtubes, which are thin long protrusions that help the tumor grow and spread. Follow-up experiments were conducted on mice in order to test what impact the tumor microtubes have on tumor regrowth after partial removal of a tumor region. The surgery was performed in isolation and along with growth-inhibiting treatments such as a tumor microtube inhibiting treatment and an anti-inflammatory treatment. Here, we propose a partial differential equation model applicable to describe the microtube driven regrowth of the cancer in the lesion. We find that the model is able to replicate the main trends seen in the experiments such as fast regrowth, larger cancer density in the lesion, and further spread into healthy tissue. The model indicates that the dominant mechanisms of re-growth are growth-inducing wound healing mechanisms. The tumor microtubes accelerate this process as the microtubes provide orientational guidance from the untreated tissue into the lesion.

## 1 Introduction

Astrocytomas are brain tumors that are extremely difficult to treat, with most treatments leading to tumor recurrence [24, 32]. Fairly recently, Osswald et al. [24] have performed mice experiments where they took grade II-IV astrocytomas from human brain tumors and implanted them into mice. By observing the growth of the astrocytoma cells, they discovered an important structure which they named “tumor microtubes.” Tumor microtubes (TMs) describe thin long protrusions extending from the cell body of an astrocytoma cell. Weil et al. [32] performed micro surgery experiments, to better understand the role of TM with respect to cancer regrowth and response to treatments. In this paper we develop a mathematical model that can describe the experiments of [32]. We find that wound healing mechanisms are likely the most important contributors to tumor regrowth, while the TM-induced anisotropy helps to accelerate this process.

### 1.1 Tumor Microtubes

TMs can extend up to 500*μ*m and the number of TMs increases as the tumor progresses [24]. Moreover, TMs have been linked to aid tumor growth, promote spatial spread and have been shown to allow communication between cells [24, 32, 14]. This is due to TMs ability to link astrocytoma cells to each other through connexin 43 gap junctions [32, 14]. These gap junctions facilitate cell communication by allowing calcium wave propagation [32, 14]. Moreover, it has been identified that the growth associated protein-43 (GAP-43) is essential for TM formation and TM network functionality [24, 32, 10]. TMs have also been linked to play a role in treatment resistance, where it was shown that the communication network between astrocytoma cells plays a significant role in aiding the repair of the tumor after treatments such as surgery, radiotherapy, or temozolomide chemotherapy [32].

In 2017, Weil et al. [32] performed detailed mice experiments in order to test astrocytoma treatments. Mice had one of the primary glioblastoma (grade (IV) astrocytoma) cell lines (S24 or T269) implanted into their brain. The main difference between these strands is that in vitro, S24 tends to form more TMs [32]. After sufficient growth of the tumor inside the mouse brain, Weil et at. [32] performed a surgical treatment where they resected a cylindrical volume from the tumor using a syringe with a diameter of approximately 300*μ*m. They examined the behaviour of the tumor for 28 days post surgery, and found that the tumor regrows faster and denser inside the lesion. In addition to surgery, they tested other treatments combined with surgery. One of the treatments they performed alongside surgery can be broadly categorized as targeted therapy, which involved specifically inactivating GAP-43 using small hairpin RNA (shGAP-43) to prevent normal formation of the TMs. They found that this was a successful treatment to delay the regrowth of the tumor in the lesion in comparison to the control treatment (surgery). Furthermore, they tested an anti-inflammatory treatment with dexamethasone (DEX) which was administered daily for 14 days, starting on the day of surgery. They found that this treatment only allowed for a transient benefit, where the tumor regrew slower on day 7 post surgery in comparison to the control treatment (surgery), but then on day 14 there was no significant difference between the control treatment (surgery) and the combination treatment (surgery combined with DEX).

Several biological processes might be responsible for the observations of fast regrowth post surgery in the mice experiments. TMs are able to aid with cell proliferation [24] and Weil et al. [32] showed that post surgery, nearby cells actually extend their TMs towards the lesion along which cell nuclei can travel. These nuclei can then begin repopulating the lesion. Weil et al. [32] also showed that astrocytoma cells can actually move towards the lesion, where on day 5 post surgery more than half astrocytoma cells move toward the lesion. In addition, surgery creates a wound and a wound healing response is triggered. Wound healing is classically a four stage process with stages that may overlap: 1) hemostasis, 2) inflammation, 3) cell proliferation, and 4) remodelling [7, 23]. Inflammation during a wound healing process has been shown to have the potential to be tumor promoting, since immune inflammatory cells release growth factors and promote cancer cell proliferation and cancer spread [13, 12]. Moreover, an injury may activate microglia and macrophages which can aid with cancer cell survival, proliferation, and invasion [21, 11]. Hence, in addition to TMs, wound healing mechanisms may also be promoting tumor growth. We will use our mathematical model to investigate the significance of these processes related to the cancer regrowth data from [32].

### 1.2 Modelling of Glioma

There have been a number of mathematical models proposed to model the growth and spread of glioblastoma. One can use agent based models to model glioblastoma dynamics to better capture single cell behaviour. For example, Gao et al. [6] have used a cellular Potts model to capture cell structure, adhesion, and motility as well as the cell heterogeneity of in a glioblastoma tumor. Reaction diffusion equations have been especially popular to use to model the growth dynamics for glioblastoma tumors, as the diffusion term can be used to capture the movement of cells and the source term (in the form of exponential or logistic growth) can be used to capture the growth dynamics of glioblastoma [22, 28]. The simulations from these models have been shown to have good agreement with MRI data of glioma tumors [22, 28] where the model in [28] can adapt to different cell movement speeds along the white or grey matter tracts in the brain and the model proposed in [22] can use Bayesian personalization of model parameters to better fit to patient data. These models can also be adapted to include anatomical boundaries [20]. However, the models in [22, 28] have only employed isotropic diffusion of the glioma cells, meaning that cells have equal probability to move in any given direction. It has been observed that glioma cells tend to move along white matter tracts in the brain, showing that glioma cells can have preferential movement [5]. To include preferential movement one can use a mechanistic model where anisotropic diffusion of cancer cells arises as a byproduct of various mechanistic forces. For example, Colombo et al. [3] proposed a fully mechanistic model which incorporated patient specific anisotropy from diffusion tensor imaging (DTI) data and accounted for nutrient availability for the cancer cells. Alternatively, a fully anisotropic version of the isotropic models described above can be formally derived in the form of transport equations [8, 25]. The anisotropic movement in the transport equation model is captured by the diffusion tensor, which can be formulated to include patient specific DTI by specifying the orientational distribution of tissue fibers [25, 4, 27].

Recently, a transport equation model was proposed by Hillen et al. [15] which incorporated not only the dynamics of glioma cells, but also the TMs. This model includes TM invasion (where the preferential movement of the tips in the anisotropic environment is accounted for), TMs ability to retract and create new TMs, nuclei movement along the TMs, nuclei maturation into glioma cells, and glioma cell division. By comparing the simulated tumor growth to the data found in Osswald et al. [24] mouse experiments, they validated the model where they showed that the model is able to replicate the trends in the data. Moreover, they showed that under certain assumptions their model can reduce to key models found in previous studies, for example the simple reaction diffusion model proposed by [28].

In this work, we propose a mathematical model applicable to the mice experiments post surgery from the experiments done by Weil et al. [32]. We greatly simplify the processes that may be occurring biologically, and derive a much simpler model than the one proposed by Hillen et al. [15]. We incorporate the directed movement of the glioblastoma cells, by assuming a radially symmetric domain and assuming that the cancer cells move radially towards the lesion. We take the base model from Bica et al. [1] to describe this radial movement. Our proposed model accounts for the wound healing process and TM dynamics within the model parameters. The model also includes the healthy cell dynamics. We find that the model is able to replicate the trends seen in the Weil et al. [32] experiments post surgery and can replicate the trends from the targeted therapy as well as the anti-inflammatory treatments. Moreover, we find that the growth advantage through TM is significant during the first 14 days of the regrowth.

## 2 The Model

Before introducing our model which will be used to recreate the experiments in Weil et al. [32], we first discuss the spatial model that we will use to describe anisotropic (and also isotropic) movement. The domain that we will work with is a two dimensional circle, which we call Ω. In this circle, the cells have the option to move along the radial lines, perpendicular to them, or simply diffuse (not have any relationship to the radial lines). This movement is sketched in Figure 1, where *α*(*r*) is a function depending on the radius, which denotes the degree of movement along the radial lines. If *α*(*r*) > 0 then the cell moves preferentially along the radial lines and if *α*(*r*) < 0 then the cell moves preferentially perpendicularly to the radial lines as shown in Figure 1. In the limits *α* → 1 cells move fully parallel to the radial lines and *α* → −1 cells move fully perpendicular to the radial lines. If *α*(*r*) = 0, then the movement is isotropic (i.e., cells diffuse with no correlation to the radial lines). As in Bica et al. [1], we assume that |*α*(*r*)| ≤ 1.

**Figure 1:**
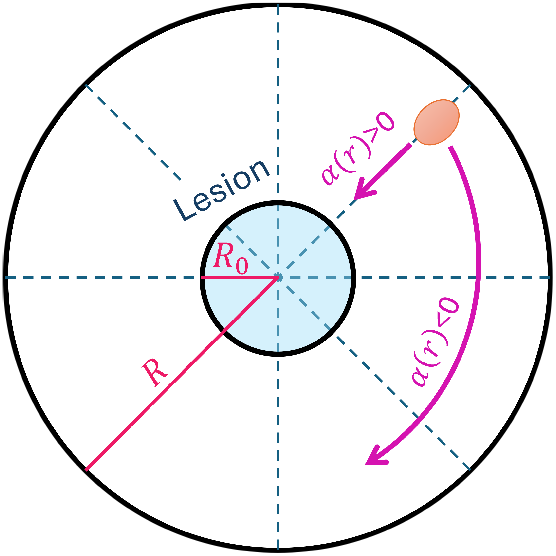
Plot of cell movement along radial lines. The orange oval denotes a cell. If *α*(*r*) > 0, then the cell moves preferentially parallel to the radial line and if *α*(*r*) < 0, then the cell moves preferentially perpendicular to the radial lines. The lesion is denoted by the filled in circle filled with blue, where *R*_0_ is its radius. The outer circle has radius *R*.

To derive the model for the directed movement shown in Figure 1, Bica et al. [1] started with the fully anisotropic diffusion equation proposed by Hillen et al. [17], which is given by

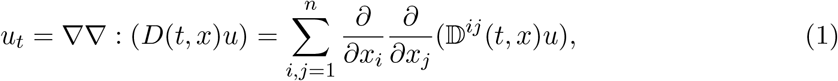

with *u* = *u*(*t, x*), *t* ∈ ℝ, *x* ∈ Ω, and where 𝔻 (*x*) is a diffusion tensor. We will transform to polar coordinates (*r, ϕ*) by assuming no-flux boundary conditions at *r* = *R*:

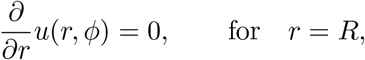

and assuming that the solutions are bounded at *r* = 0.

The diffusion tensor used by Bica et al. [1] was derived by Hillen et al. [18] from the von Mises distribution, and the formula for the diffusion tensor is given by

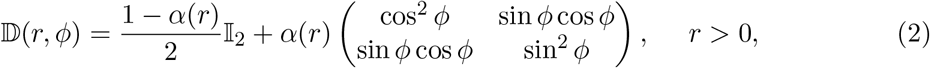

in the planar polar coordinate system (*r, ϕ*). In (2), *α*(*r*) is the function describing the degree to which cells move along the radial lines as discussed earlier. Substituting (2) into (1), Bica et al. [1] derived the following model

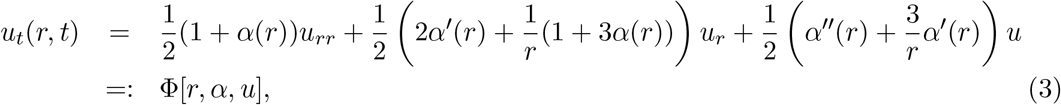

where *r* ∈ (0, *R*] with *R* being the maximum radius, and Φ[*r, α, u*] denotes the linear second order differential operator that depends on *r* and *α*(*r*). Note that in the isotropic case *α* = 0, this operator simplifies to the standard Laplacian in polar coordinates in the radially symmetric case with diffusion coefficient 1/2:

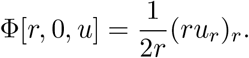

The model (3) is a one dimensional PDE model which is applicable to situations in Figure 1 and can be easily solved numerically.

To describe the experiments of Weil et al. [32] we extend the spatial model for two competing populations, cancer cells and healthy tissue. Let *u*(*t, r*) ≥ 0 be the density of the glioma cells and *v*(*t, r*) ≥ 0 be the density of healthy cells, where *r* > 0. The model we propose is as follows

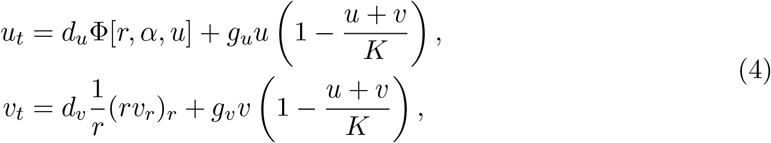

where Φ is given in (3).

In (4), *α*(*r*) is function describing the degree to which cells move along the radial lines as discussed earlier. The parameters *d*_*u*_ and *d*_*v*_ are the respective diffusion coefficients of cancer cells and healthy cells, *g*_*u*_ and *g*_*v*_ are the respective growth rates of cancer cells and healthy cells, and *K* is the carrying capacity. Notice that in the second equation, the spatial term is obtained by setting *α*(*r*) = 0, i.e. using Φ[*r*, 0, *v*]. Essentially, in Model (4), we are assuming that glioblastoma cells can move along the radial lines and the healthy cells simply move by diffusing. Both the cancer and healthy cells grow logistically where both cell types compete for the limited space and nutrients which is capped at the carrying capacity. Note that in Model (4), we do not include the dynamics from the wound healing, and we will instead incorporate it in the parameters later.

### 2.1 Parameterization

We take our domain Ω to be a circle with a radius of *R* = 600*μ*m. The domain Ω will have two distinct regions: inside the lesion (including the lesion boundary) and the region outside the lesion (see Figure 1). We call the lesion and its boundary Ω_*in*_ and the region outside the lesion Ω_*out*_. The radius for Ω_*in*_ is taken to be *R*_0_ = 150*μ*m since the diameter of the lesion in Weil et al. [32] Figure 1 is approximately 300*μ*m. We assume that the interiors of Ω_*in*_ and Ω_*out*_ are homogeneous. Ω_*out*_ contains blood vessels and other tissue structures, but they are not the focus of this study and we assume they are essentially uniformly distributed.

Due to the TMs and wound healing mechanisms, the cells growing in the lesion area will have an advantage, therefore requiring us to distinguish between parameters in Ω_*in*_ and in Ω_*out*_. Hence, to summarize the parameters, we represent them in a tuple (· ; ·) in Table 1, where the first entry is the parameter inside Ω_*in*_ and the second entry is inside Ω_*out*_. To make it clear which parameter we discuss, we use the notation 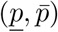 where <u>*p*</u> denotes the parameter inside Ω_*in*_ and 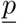 denotes the parameter in Ω_*out*_.

**Table 1:** Summary of the parameters used. The tuple denotes parameters within and outside the lesion. The first value contains the parameter inside the lesion, and the second value contains the parameter outside the lesion.

| Units | $\mu\text{m}^2/\text{h}$ | $\mu\text{m}^2/\text{h}$ | $1/\text{h}$ | $1/\text{h}$ | $\text{cells}/\mu\text{m}^2$ |
| --- | --- | --- | --- | --- | --- |
| Parameter | $(\underline{d}_u, \bar{d}_u)$ | $(\underline{d}_v, \bar{d}_v)$ | $(\underline{g}_u, \bar{g}_u)$ | $(\underline{g}_v, \bar{g}_v)$ | $(\underline{K}, \bar{K})$ |
| Value | (6.4; 6.4) | (3.2; 1.6) | (0.018; 0.006) | (0.0012; 0.0006) | (0.01, 0.0018) |

In Table 1, we choose to set the diffusion coefficients 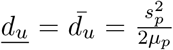 as was done in [15] where the factor of 1*/*2 comes from (3). This estimate is based on a typical cell speed of *s*_*p*_ = 4.8*μ*/h and a turning rate of *μ*_*p*_ = 1.8/h, which are parameters taken from [15]. For *d*_*v*_, we assume that inside Ω_*in*_, <u>*d*_*u*_</u> = <u>*d*_*v*_</u>/2, and inside 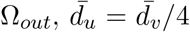, because healthy cells move slower than cancer cells [13], and since there is more space available in Ω_*in*_ the cells can move faster. To obtain *g*_*u*_ for cancer cells, we do a rough estimate based on the figures given in Weil et al. [32] which we summarize here. From Figure 1 in [32] (also shown in Figure 3, bottom two rows), we see that the rough doubling time in Ω_*out*_ is 5 days based on images of day 3 and day 5 in [32]. Then using the formula growth rate = *ln*(2)*/*doubling time [19], we get a rough growth rate of 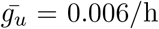 in Ω_*out*_. To get the growth rate for cancer cells in Ω_*in*_, we triple the 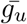 from Ω_*out*_ as cancer cells grow faster inside the lesion due to the wound healing and TMs. For the healthy cell growth rate, we assume that 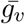 is the same as the bulk growth rate of a glioma tumor, hence we set it to be 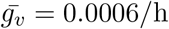 h [15]. Since the cells in Ω_*in*_ have an growth advantage due to the wound healing response, we double the growth rate in Ω_*in*_ and obtain that 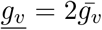. We also estimate the carrying capacity from Figure 1 in [32]. For 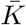 in Ω_*out*_, based on day 0 in Figure 1 in [32], we can roughly count how many cancer cells fit into a 150*μ*m × 150*μ*m square. We then take that estimate as 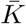 for Ω_*out*_ for both healthy and cancer cells. For <u>*K*</u> in Ω_*in*_, we note that a glioma cell is approximately 10*μ*m from Figure 2 in [32] and estimate that 225 cells fit into a 150*μ*m × 150*μ*m square. From there we obtain *K* in Ω_*in*_. Note that we assume a higher carrying capacity in Ω_*in*_ due to wound healing signalling and angiogenesis hence cells in Ω_*in*_ have more favourable growth conditions [9].

**Figure 2:**
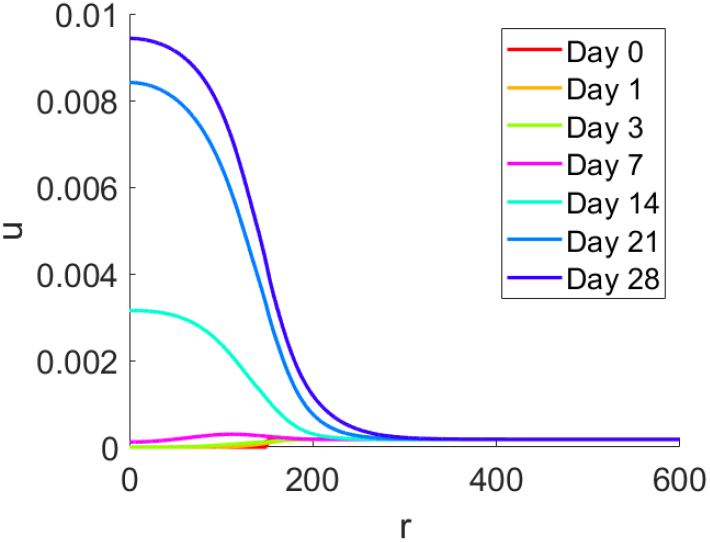
Plots of 1D solutions of Model (4) at particular days which illustrate the cancer cell density *u* with respect to the radius *r*. Model (4) was simulated with parameters given in Table 1, initial conditions (6), and boundary conditions (7).

**Figure 3:**
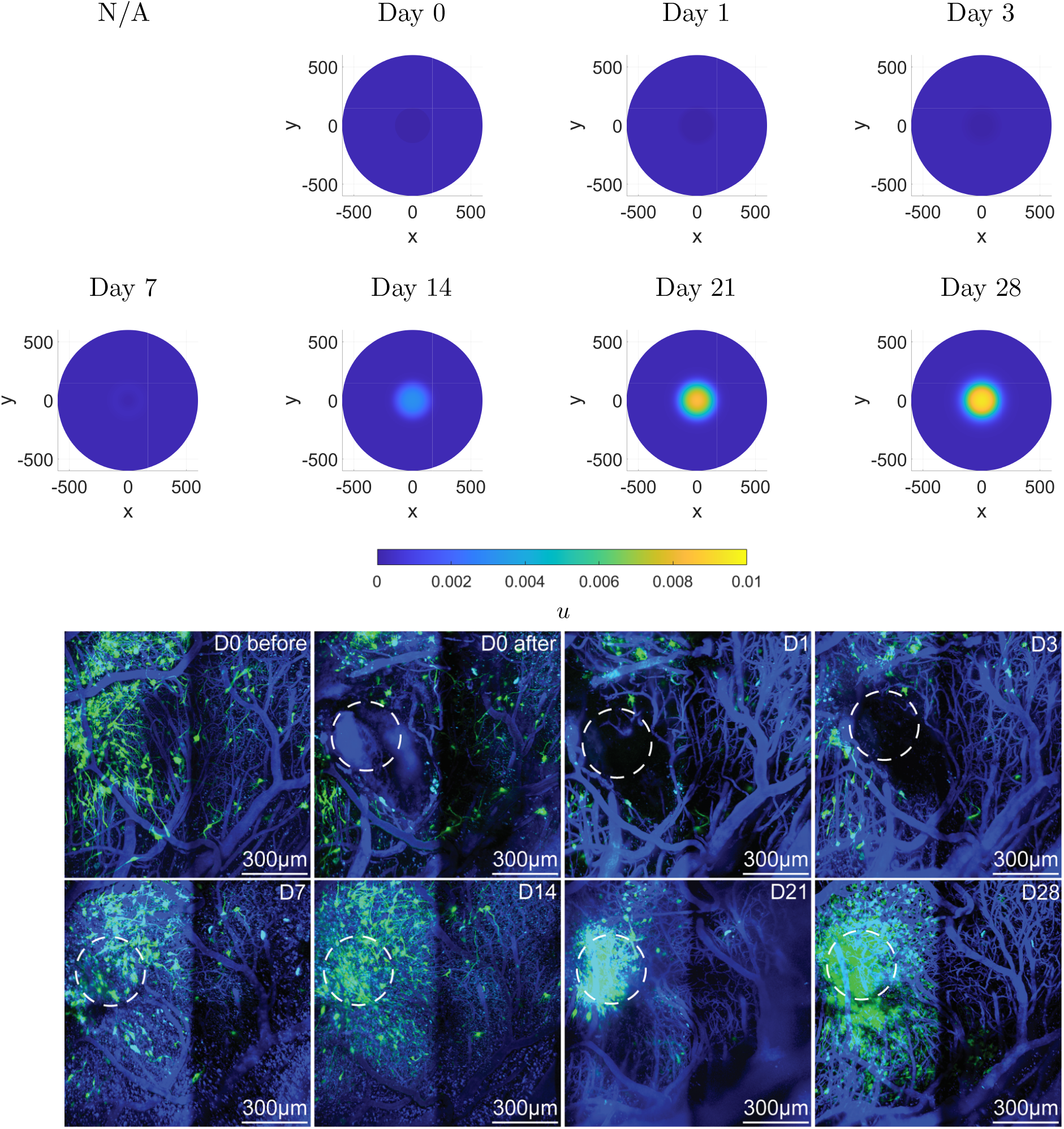
Comparison of the simulated tumor density (first two rows) with experimental results (last two rows). The first two rows show plots of 2D solutions of Model (4) at particular days where each plot illustrates the cancer cell density *u* on a circle Ω. The radius of Ω is 600*μ*m. Model (4) was simulated with parameters given in Table (1), initial conditions (6), and boundary conditions (7). The last two rows show the experimental images of gliomblastoma growth from Weil et al. [32]. The dashed circle is the lesion cite, in green are the glioblastoma cells, and in blue are the blood vessels (used with permission).

For the function *α*(*r*) we choose a hyperbolic tangent function given by

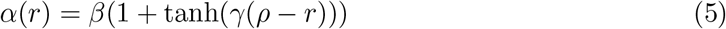

which is one of the functions for *α*(*r*) proposed in [1]. We set *ρ* = 150 because between the regions *r* ∈ (0, 150] (Ω_*in*_) and *r* ∈ (150, 600] (Ω_*out*_), the degree of anisotropy changes. We reason that the closer the cells are to the lesion, the more radially they move as evidence from [32]. In [32], they note that nearby cancer cells tend to move toward the lesion. Hence, we set *β* = 0.3 which makes the upper bound of *α*(*r*) to be 0.6, meaning that most cells inside and near Ω_*in*_ move parallel to the radial lines. As the cells are further away from the lesion, we assume they mainly move via diffusion, so *α*(*r*) → 0 as *r* → *R*. We set *γ* = 0.06 for a smooth transition in *α*(*r*) between 0.6 and 0. We could also choose *γ* = 1 for a sharper transition. We summarize the parameters choices for *α*(*r*) in Table 2 and plot the two possible choices of *α*(*r*). Note that for the isotropic case we have *α*(*r*) = 0.

**Table 2:** Summary of parameters used for *α*(*r*) (left). Plot of the smooth and sharp cases for *α*(*r*) (right).

| $\alpha(r)$ type | $\beta$ | $\gamma$ | $\rho$ |
| --- | --- | --- | --- |
| Smooth | 0.3 | 0.06 | 150 |
| Sharp | 0.3 | 1 | 150 |
| Isotropic | 0 | 0 | 0 |

For model (4), we use the following initial condition,

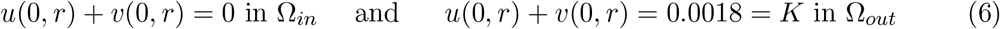

meaning that there are no cells in the lesion, and outside the lesion the cell density is at carrying capacity. We assume that the cancer cells make up 10% of the total cell density outside the lesion. The boundary conditions for model (4) are Neumann no flux boundary conditions. That is

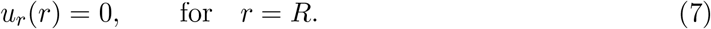

## 3 Results

### 3.1 Simulation of the Model

To simulate Model (4) with the initial conditions (6) and boundary conditions (7), we use Matlab’s pdepe solver. Using the parameters in Table 1 and Table 2 applicable to the “smooth” case, we obtain the 1D solutions for Days 0, 1, 3, 7, 14, 21, 28 which are illustrated in Figure 2. The solutions in Figure 2 were then rotated radially to obtain the visualization in Ω as shown in the first two rows of Figure 3.

In Figure 2, we see that the cancer density *u* invades the lesion region Ω_*in*_ as time progresses and by Day 28, *u* almost grows to the carrying capacity *K* = 0.01cells/*μ*m^2^ which is also illustrated in Figure (3). We chose to visualize these particular days in order to see if we could match the trends seen in Weil et al. [32] Figure 1 (bottom two rows of Figure 3). Indeed, with these chosen parameters we are able to replicate the general growth speed and general shape of the tumor. In Figure 3 from the bottom two rows it can be seen visually that on day 7 the tumor has started regrowing visually matching the density on day 0. Here, we see a similar effect where on day 7 (in the top two rows), the density inside the lesion starts to match the density outside the lesion. In our simulations, we see that the tumor has grown significantly inside the lesion on day 14, which matches the observations in [32]. We note that on this day, the tumor starts growing outside the lesioned area as well. On day 21, the tumor has more than doubled in size, which can also be seen in [32]. The tumor keeps growing in size on day 28, almost reaching the carrying capacity *K* = 0.01cells/*μ*m^2^ inside the lesion. On days 14, 21 and 28, we see from Figure 2 that the cancer *u* not only grows inside the lesion, but outside the lesion as well, where *u* progressively invades the nearby space of the lesioned area. Evidence of this is also shown in [32], where the tumor begins growing outside the lesion as well, which is particularly evident on day 21 and onward.

From the above observations, we believe that Model (4), does well in replicating the qualitative observations seen in [32]. However, due to a lack of quantitative measures, such as the *u* density per *μ*m^2^, we cannot validate the model further.

### 3.2 Isotropic and anisotropic diffusion

In this subsection, we study how the degree of anisotropy affects the model. As discussed before, setting *α*(*r*) = 0 and its consecutive derivatives to zero in (3) yields a model with isotropic diffusion. We will study if this causes any significant changes to the dynamics that are seen in Figure (3). Further, it is of interest to examine how the degree of smoothness in the *α*(*r*) affects the dynamics. We will study this by simulating model (4) for each choice of *α*(*r*) in Table 2.

In Figure 4, we see a comparison of what occurs to the cancer cell density *u* for different choices of *α*(*r*) at particular days 3, 14, and 28. For each case, we use the parameters in Table 1 and parameters for *α*(*r*) vary based on the case as listed in Table 2.

**Figure 4:**
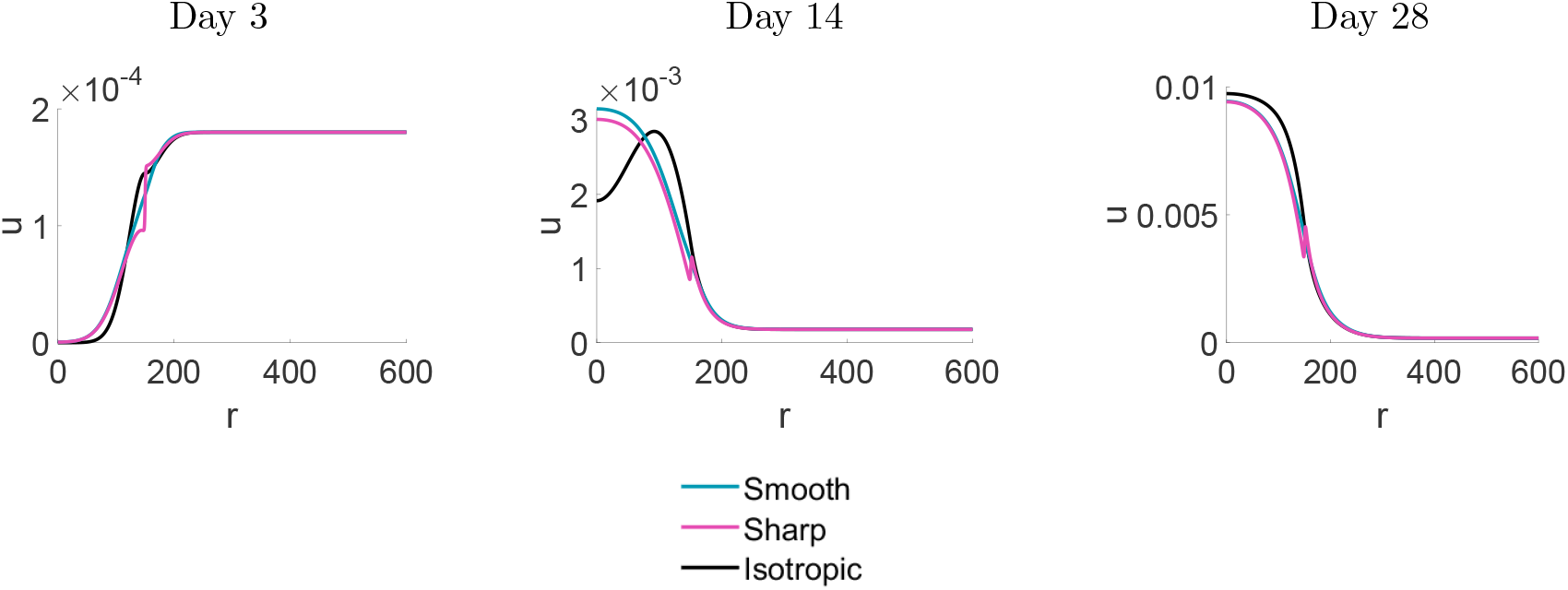
Plots 1D solutions of Model (4) at particular days which illustrate the cancer cell density *u* for each *α*(*r*) choice with respect to the radius. Model (4) was simulated with parameters given in Table (1), initial conditions (6), and boundary conditions (7).

The blue curves in Figure 4 refers to the sharp *α*(*r*) case, the pink curves refer to smooth *α*(*r*) case and the black curves refer to the isotropic case (*α*(*r*) ≡ 0). We see that the day 3 dynamics for *u* are very similar in each case, with only slight differences. The sharp *α*(*r*) case has sharper transitions between the cancer density within the lesion and the cancer cell density outside the lesion whereas the transition for the smooth *α*(*r*) case is smooth between the regions. We see that the anisotropic cases (smooth and sharp *α*(*r*)) show that the cancer cell density invades further into the lesion initially than the isotropic case (seen by the leading edge on the left). The isotropic case shows steady and slower invasion into the lesion, which is also seen by the peak in the black curve on day 14. On day 14, we see that the isotropic cases have completely invaded the interior of the lesion while the isotropic case is still growing. On day 28, there is little difference between the cases.

Further, for each case (isotropic and anisotropic), we see that the cancer cells invade the healthy tissue outside the lesion at very similar speeds. This overgrowth was also observed in the experiments [32].

For the sharp case, we see in the pink profiles in Figure 4 that a local peak forms near the boundary of the lesion at *r* = 150. This peak would yield a ring around the bulk tumor if we were to plot these cases in 2D. The presence of this peak is, indeed, not surprising, as it was shown that the anisotropic diffusion equation (1), with sharp transitions in the diffusion coefficient, can form singularities at such transition layers [16]. Such a ring is not observed in the experiments of [32].

### 3.3 Sensitivity

In addition, we study what happens to the dynamics for each case (smooth, sharp and isotropic *α*(*r*)) if the parameters in Table 1 vary. The results are summarized in Table 3. We choose to examine the most interesting variations, which are those that may affect the growth of *u*. For all three cases, if the carrying capacity <u>*K*</u> is lowered but does not decrease beyond 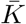 , then there is no effect on the speed of growth of *u*. Note, that this trend still holds if *K* is increased. If the growth rate of cancer <u>*g*_*u*_</u> is lowered but not lower than 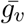, then *u* grows slower, as expected. Similarly, if <u>*g*_*u*_</u> is increased then *u* grows faster in all three cases. If we set the growth rates to be the same for both the cancer cell and healthy cell densities, then *u* will not grow in all three cases. In this situation, the healthy cell density *v* is the dominant density in both regions (Ω_*in*_ and Ω_*out*_). Finally, increasing the diffusion of *v*, does not impact the growth of *u* significantly, in all three cases. Note, that for this study, the changes in parameters were significant, and not small perturbations that we will do when performing the sensitivity analysis in the following section. We did this analysis to see if there were any significant changes in the dynamics between the cases when varying parameters. After performing this analysis, we see that the three cases (smooth, sharp, and isotropic *α*(*r*)) have very similar overall dynamics, where the anisotropy accelerates the regrowth initially.

**Table 3:** Summary of trends from varying parameters for the three cases of *α*(*r*) (smooth, sharp, isotropic). Note that when we write 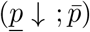 we do not decrease the parameter past 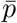 , that is, 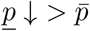.

| $\alpha(r)$ | $(\underline{d}_u; \bar{d}_u)$ | $(\underline{d}_v; \bar{d}_v)$ | $(\underline{g}_u; \bar{g}_u)$ | $(\underline{g}_v; \bar{g}_v)$ | $(\underline{K}; \bar{K})$ | Result in $\Omega_{in}$ |
| --- | --- | --- | --- | --- | --- | --- |
| Isotropic | - | - | - | - | $(\underline{K} \downarrow; \bar{K})$ | $u$ grow at the same pace |
| Isotropic | - | - | $(\underline{g}_u \downarrow; \bar{g}_u)$ | - | - | $u$ grows much slower |
| Isotropic | - | - | $(\bar{g}_v; \bar{g}_v)$ | $(\bar{g}_v; \bar{g}_v)$ | - | $u$ does not grow |
| Isotropic | - | $(\underline{d}_v \uparrow; \bar{d}_v \uparrow)$ | - | - | - | $u$ grow at the same pace |
| Smooth | - | - | - | - | $(\underline{K} \downarrow; \bar{K})$ | $u$ grow at the same pace |
| Smooth | - | - | $(\underline{g}_u \downarrow; \bar{g}_u)$ | - | - | $u$ grows much slower |
| Smooth | - | - | $(\bar{g}_v; \bar{g}_v)$ | $(\bar{g}_v; \bar{g}_v)$ | - | $u$ does not grow |
| Smooth | - | $(\underline{d}_v \uparrow; \bar{d}_v \uparrow)$ | - | - | - | $u$ grow at the same pace |
| Sharp | - | - | - | - | $(\underline{K} \downarrow; \bar{K})$ | $u$ grow at the same pace |
| Sharp | - | - | $(\underline{g}_u \downarrow; \bar{g}_u)$ | - | - | $u$ grows much slower |
| Sharp | - | - | $(\bar{g}_v; \bar{g}_v)$ | $(\bar{g}_v; \bar{g}_v)$ | - | $u$ does not grow |
| Sharp | - | $(\underline{d}_v \uparrow; \bar{d}_v \uparrow)$ | - | - | - | $u$ grow at the same pace |

### 3.4 Sensitivity analysis

We perform a sensitivity analysis following the procedure outlined in [31]. Note, that we adapt the sensitivity analysis for ODEs to PDEs by fixing the spatial point inside the lesion (*r* = 20).

The sensitivity indices 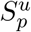 for the cancer cell density *u* and given parameter *p* is summarized in Figure 5. The sensitivity indices were calculated numerically for a fixed location inside the lesion *r* = 20*μ*m at time 28 days according to the relative change [31]

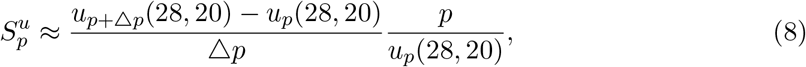

where *p* is the parameter of interest, △*p* is 1% of the *p* value, *u*_*p*_(28, 20) is the solution with default parameters *p* at location *r* = 20, *t* = 28, and *u*_*p*+△*p*_ is the solution with a slightly increased parameter value. We see in Figure 5 that the parameters that have the greatest impact in affecting the cancer cell density are the cancer growth rate *g*_*u*_ and carrying capacity *K*. This is not surprising as increasing growth rate allows for the cancer cell density to grow faster, and increasing the carrying capacity allows for the cancer cell density to grow larger. Increasing the growth rate of the healthy cell density *g*_*u*_ and increasing the diffusion constant of the healthy cell density *d*_*u*_ have minor negative effect on the growth of the cancer cell density. This is as expected, since faster growing healthy cell density can populate more space decreasing the cancer cell density, and increasing the diffusion constant allows for the healthy cell density to spread more, thereby decreasing space available for the cancer cells. The sensitivity analysis reveals that increasing the degree of anisotropy (increasing *β* in *α*(*r*)) slightly decreases the growth of the cancer cell density. This means that moving via isotropic diffusion is slightly more advantageous in increasing cancer cell density. This is what we see in Figure 4, where the isotropic movement of cancer cell density has a more concentrated core. Negative sensitivity to *d*_*u*_ can be explained by noting that higher diffusion will cause more cancer cells to move out from the lesion area Ω_*in*_ and those cells lose the additional growth benefit Ω_*in*_ provides.

**Figure 5:**
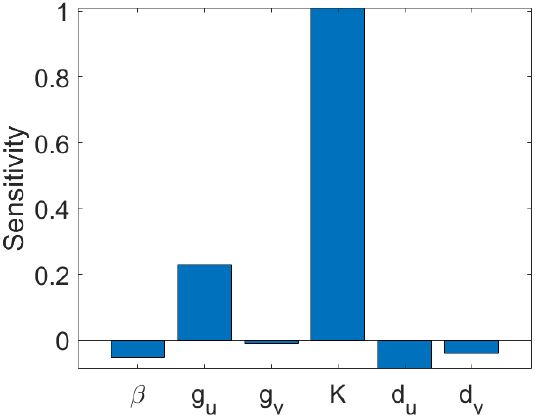
Summary of the sensitivity indices for the cancer cell density *u*. The parameters in the image refer to the parameters in Ω_*in*_.

## 4 Treatment

Here, we test two treatments in combination with surgery as was done in [32]. The first is a targeted therapy which was performed by injecting a small hairpin RNA (shGAP-43) to inhibit TM formation (by inhibiting GAP-43). We apply this treatment by reducing the growth rate of cancer cells by 50% in the lesion, and we reduce the value of *α*(*r*) by 75% in Ω_*in*_ because targeted therapy targets the TMs reducing their ability to form, hence hindering directed motion and cancer cell proliferation. We see the results of this treatment in Figure 6 middle row. Note that the colorbar differs from that of Figure 3, in order to see the smaller densities easier. For the targeted therapy treatment, we see that the cancer cell density grows significantly slower than the control. The same effect was seen in Figure 3 in [32] and we are able to recreate the dynamics for this treatment well.

**Figure 6:**
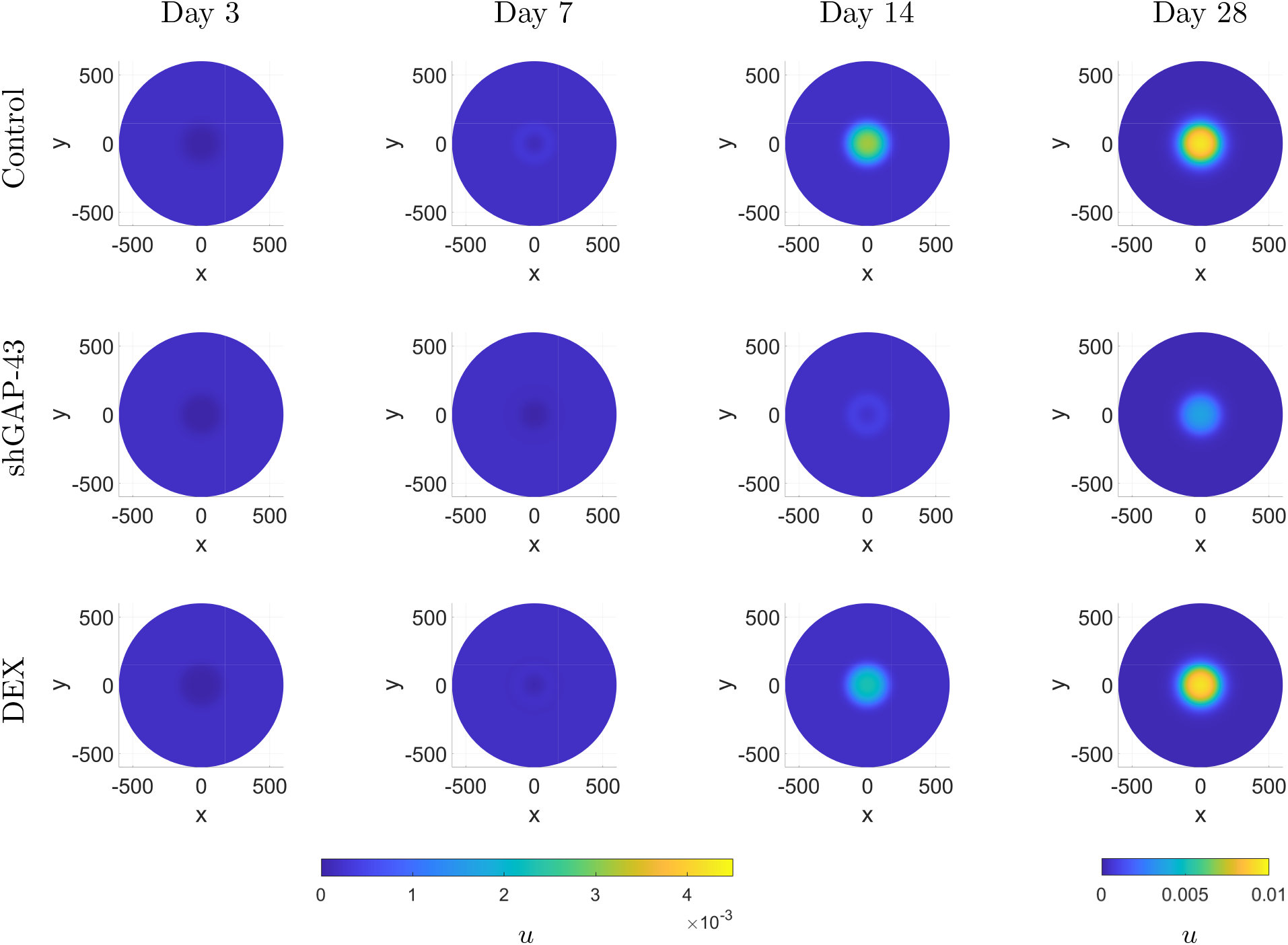
Illustration of different treatments where the colorbar shows the cancer cell density *u*. In row 1, Model (4) is simulated with the control parameters in Table 1. In row 2, targeted therapy with shGAP-43 is simulated (4) and in row 3, the anti-inflammation treatment is simulated by using (4) and the parameters outlined in Section 4. For days 3, 7, 14 the colorbar on the left applies and for day 28 the colorbar on the right applies in order to show changes in the densities more clearly.

The other treatment we test is the anti-inflammatory treatment with dexamethasone (DEX) [32]. DEX is a glucocorticoid (GC) which binds to the GC-receptor (GR) of the cells and deactivates NF-*κ*B [29, 2, 33]. NF-*κ*B is essential for the homeostasis of the immune system and lower levels of NF-*κ*B lead to reduced immune response [2]. However, the immune suppression is not permanent and several resistance mechanism are known [30, 26]. For example in leukemia it was shown [30] that immune cells exposed to GC reduce the expression of GR over time, consequently becoming less sensitive to glucocorticoids.

We employ the following sigmoidal curve to describe how the model parameters return to their base values as treatment resistance sets in. We define

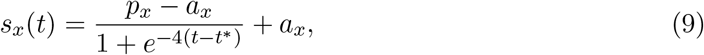

where the sub-index *x* stands for either of *x* = *g*_*u*_, *g*_*v*_, *K*, i.e. the two growth rates and the carrying capacity. Here *p*_*x*_ denotes the base value in Table (1), *a*_*x*_ is the value of the parameter under DEX treatment. The time *t*^∗^ = 3.5 is set to describe onset of resistance between 3 and 4 days, which was the best choice to describe the experimental observations. In *s*_*K*_(*t*), we set 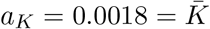 while *p*_*K*_ = 0.01 = <u>*K*</u>. For the growth rates *g*_*u*_ and *g*_*v*_ we assume that both are initially reduced by 50% inside the lesion. Hence *a*_*gu*_ = <u>*g*_*u*_</u>/2, *p* = <u>*g*_*u*_</u> and *a* = <u>*g*_*v*_</u>/2, *p* = <u>*g*_*v*_</u> with *g*_*u*_ and *g*_*v*_ from Table (1).

The model we use inside Ω_*in*_ during the DEX treatment is

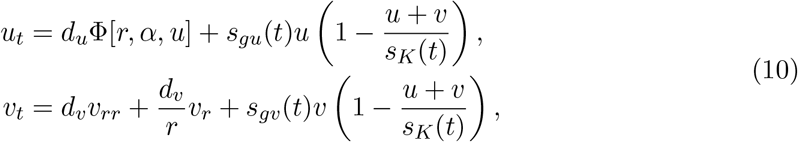

whereas model (4) is unchanged in Ω_*out*_. Note that (9) could be replaced by any other sigmoid function with very similar results.

The simulations for the DEX treatment are shown in the third row of Figure 6. We can see that the cancer cell density is lower on Day 7 than in the control, and then on day 14 it begins to catch up and by day 28, it is at the same level as the control. In the corresponding Figure 4 in [32] they report that there is no significant difference between the control and the DEX treated tumor after 14 days or more.

## 5 Discussion

In this study, we have proposed a mathematical model that applies to the mice experiments conducted by Weil et al. [32] post surgery of astrocytoma tumors. We find that the model is able to replicate the dynamics of glioma tumor growth and spread post surgery. Our model heavily simplifies the wound healing mechanisms as well as TM dynamics, which serve to elevate the proliferation rates of cells and carrying capacity within the lesion site. The simulations show that the tumor grows back faster and denser within the lesion site matching the observations seen in Weil et al. [32]. This shows that the wound healing processes and the proliferative advantage from the TMs are potentially the key to explaining the faster and denser regrowth that was observed experimentally.

Our model is based on an anisotropic model proposed by Bica et al. [1], in order to account for the potential directed motion of astrocytoma cells. We found that anisotropic movement speeds up the regrowth as compared to the isotropic case. However, with the available measurements we find that isotropic as well as anisotropic diffusion can explain the data well.

We tested the combined treatments: surgery with targeted therapy (shGAP-43) and surgery with an anti-inflammatory treatment (DEX). Due to a lack of data, we kept the addition into the model simple where treatments predominately affect the growth and carrying capacity parameters. Our model was able to generally match the trends seen in the treatment experiments of Weil et al. [32], where the targeted therapy matches the experimental data well but the anti-inflammatory treatment is not quite able to match the speed of regrowth that is seen in the experiments. We find that the treatments that are able to significantly reduce the proliferation rate of cancer cells as well as the carrying capacity for a prolonged period of time, are most effective at delaying the growth of the tumor. The targeted therapy treatment performs much better than the anti-inflammatory treatment as it is able to significantly reduce the proliferation rate of cancer cells by inhibiting the TMs whereas the anti-inflammatory treatment only provides a transient benefit.

We note that Model (4) can be extended to include more aggressive competition between cell types, as follows

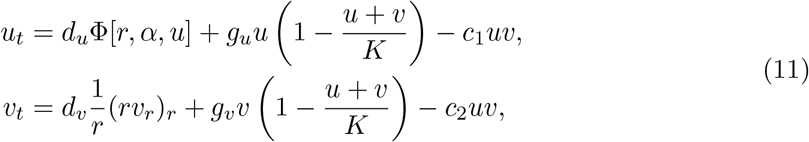

where *c*_1_ is the fitness rate of healthy cells and *c*_2_ is the fitness rate of cancer cells. That is, if *c*_1_ is higher than *c*_2_, then healthy cells are more fit than the cancer cells therefore more easily grow in the domain (and vise versa). We analyzed Model (11) but we found that Model (4) is sufficient at explaining the results of Weil et al. [32] experiments.

Our study has several limitations. We assumed that the domain outside the lesion site is homogeneous, where blood vessels are ignored. It is possible that the cancer cells would prefer to grow near the blood vessels to have easier access to nutrients which could explain the asymmetrical tumors that are seen in the experimental images. Further, we did not include a full immune response into the model. In order to obtain biologically realistic parameters for the model extension, more data needs to be collected, in particular the movement and activity of immune cells. If more data is available, more accurate effects of treatments could be incorporated into the model. Further, more complicated interactions between healthy and cancer cells could be studied, such as competition for nutrients. We refrained from putting in extra processes in order to keep the model simple and to study the main processes.

## Acknowledgments

AS acknowledges the funding from the Alberta Graduate Excellence Scholarship. TH acknowledges support from the Natural Sciences and Engineering Research Council of Canada (NSERC).

## Notes

### Competing Interest Statement

The authors have declared no competing interest.

## References

[1] Bica, I., Hillen, T., and Painter, K. J. Aggregation of biological particles under radial directional guidance. Journal of Theoretical Biology 427 (2017), 77–89.

[2] Bosscher, K. D., Berghe, W. V., Vermeulen, L., Plaisance, S., Boone, E., and Haegeman, G. Glucocorticoids repress nf-κb-driven genes by disturbing the interaction of p65 with the basal transcription machinery, irrespective of coactivator levels in the cell. PNAS 97, 8 (2000), 3919–39924.

[3] Colombo, M. C., Giverso, C., Faggiano, E., Boffano, C., Acerbi, F., and Ciarletta, P. Towards the personalized treatment of glioblastoma: integrating patient-specific clinical data in a continuous mechanical model. PLoS One 10, 7 (2015), e0132887.

[4] Conte, M., and Surulescu, C. Mathematical modeling of glioma invasion: acid- and vasculature mediated go-or-grow dichotomy and the influence of tissue anisotropy. Applied Mathematics and Computation 407 (2021), 126305.

[5] G. Gritsenko, P., Ilina, O., and Friedl, P. Interstitial guidance of cancer invasion. The Journal of Pathology 226, 2 (2012), 185–199.

[6] Gao, X., McDonald, J. T., Hlatky, L., and Enderling, H. Acute and fractionated irradiation differentially modulate glioma stem cell division kinetics. Cancer Research 73, 5 (2013), 1481–1490.

[7] Garraud, O., Hozzein, W. N., and Badr, G. Wound healing: time to look for intelligent,’natural’immunological approaches? BMC Immunology 18 (2017), 1–8.

[8] Gholami, A., Mang, A., and Biros, G. An inverse problem formulation for parameter estimation of a reaction–diffusion model of low grade gliomas. Journal of Mathematical Biology 72 (2016), 409–433.

[9] Giulian, D., Chen, J., Ingeman, J., George, J., and Noponen, M. The role of mononuclear phagocytes in wound healing after traumatic injury to adult mammalian brain. Journal of Neuroscience 9, 12 (1989), 4416–4429.

[10] Goslin, K., Schreyer, D. J., Skene, J. P., and Banker, G. Development of neuronal polarity: Gap-43 distinguishes axonal from dendritic growth cones. Nature 336, 6200 (1988), 672–674.

[11] Hambardzumyan, D., Gutmann, D. H., and Kettenmann, H. The role of microglia and macrophages in glioma maintenance and progression. Nature Neuroscience 19, 1 (2016), 20–27.

[12] Hanahan, D. Hallmarks of cancer: new dimensions. Cancer Discovery 12, 1 (2022), 31–46.

[13] Hanahan, D., and Weinberg, R. A. Hallmarks of cancer: the next generation. Cell 144, 5 (2011), 646–674.

[14] Hausmann, D., Hoffmann, D. C., Venkataramani, V., Jung, E., Horschitz, S., Tetzlaff, S. K., et al. Autonomous rhythmic activity in glioma networks drives brain tumour growth. Nature 613, 7942 (2023), 179–186.

[15] Hillen, T., Loy, N., Painter, K. J., and Thiessen, R. Modelling microtube driven invasion of glioma. Journal of Mathematical Biology 88, 1 (2024), 4.

[16] Hillen, T., Painter, K., and Winkler, M. Anisotropic diffusion in oriented environments can lead to singularity formation. European J. Applied Math. (2012).

[17] Hillen, T., and Painter, K. J. Transport and anisotropic diffusion models for movement in oriented habitats. In Dispersal, individual movement and spatial ecology: A mathematical perspective. Springer, 2013, pp. 177–222.

[18] Hillen, T., Painter, K. J., Swan, A. C., and Murtha, A. D. Moments of von Mises and Fisher distributions and applications. Mathematical Biosciences and Engineering 14, 3 (2017), 673–694.

[19] Iworima, D. G., Baker, R. K., Ellis, C., Sherwood, C., Zhan, L., Rezania, A., et al. Metabolic switching, growth kinetics and cell yields in the scalable manufacture of stem cell-derived insulin-producing cells. Stem Cell Research & Therapy 15, 1 (2024), 1.

[20] Jacobs, J., Rockne, R. C., Hawkins-Daarud, A. J., Jackson, P. R., Johnston, S. K., Kinahan, P., et al. Improved model prediction of glioma growth utilizing tissue-specific boundary effects. Mathematical Biosciences 312 (2019), 59–66.

[21] Juliano, J., Gil, O., Hawkins-Daarud, A., Noticewala, S., Rockne, R. C., Gallaher, J., et al. Comparative dynamics of microglial and glioma cell motility at the infiltrative margin of brain tumours. Journal of The Royal Society Interface 15, 139 (2018), 20170582.

[22] Lê, M., Delingette, H., Kalpathy-Cramer, J., Gerstner, E. R., Batchelor, T., Unkelbach, J., et al. Bayesian personalization of brain tumor growth model. In Medical Image Computing and Computer-Assisted Intervention–MICCAI 2015: 18th International Conference, Munich, Germany, October 5-9, 2015, Proceedings, Part II 18 (2015), Springer, pp. 424–432.

[23] MacKay, D., and Miller, A. L. Nutritional support for wound healing. Alternative Medicine Review 8, 4 (2003).

[24] Osswald, M., Jung, E., Sahm, F., Solecki, G., Venkataramani, V., Blaes, J., et al. Brain tumour cells interconnect to a functional and resistant network. Nature 528, 7580 (2015), 93–98.

[25] Painter, K., and Hillen, T. Mathematical modelling of glioma growth: the use of diffusion tensor imaging (dti) data to predict the anisotropic pathways of cancer invasion. Journal of Theoretical Biology 323 (2013), 25–39.

[26] Rosenberg, A. S. From mechanism to resistance – changes in the use of dexamethasone in the treatment of multiple myeloma. Leukemia & Lymphoma 64, 2 (2023), 283–291. PMID: 36308022.

[27] Swan, A., Hillen, T., Bowman, J. C., and Murtha, A. D. A patient-specific anisotropic diffusion model for brain tumour spread. Bulletin of Mathematical Biology 80 (2018), 1259–1291.

[28] Swanson, K. R., Alvord Jr, E. C., and Murray, J. D. A quantitative model for differential motility of gliomas in grey and white matter. Cell Proliferation 33, 5 (2000), 317–329.

[29] Tsao, P. W., Suzuki, T., Totsuka, R., Murata, T., Takagi, T., Ohmachi, Y., et al. The effect of dexamethasone on the expression of activated nf-κb in adjuvant arthritis. Clinical Immunology and Immunopathology 83, 2 (1997), 173–178.

[30] Wandler, A., Huang, B., et al. Loss of glucocorticoid receptor expression mediates in vivo dexamethasone resistance in t-cell acute lymphoblastic leukemia. Leukemia 34, 8 (2020), 2025–2037.

[31] Wang, H. Mathematical modeling I-preliminary. Bookboon, 2012.

[32] Weil, S., Osswald, M., Solecki, G., Grosch, J., Jung, E., Lemke, D., et al. Tumor microtubes convey resistance to surgical lesions and chemotherapy in gliomas. Neuro-Oncology 19, 10 (2017), 1316–1326.

[33] Yakimchuk, K. Mathematical modeling of immune modulation by glucocorticoids. Biosystems 187 (2020), 104066.

